# Surface passivation and functionalization for interferometric scattering microscopy

**DOI:** 10.1101/2024.03.30.587437

**Authors:** Jenny Sülzle, Laila Elfeky, Suliana Manley

**Affiliations:** Institute of Physics and Institute of Bioengineering, Laboratory of Experimental Biophysics (LEB), École Polytechnique Fédérale de Lausanne (EPFL), Switzerland

**Keywords:** surface passivation, surface functionalization, surface immobilization, interferometric scattering, label-free microscopy

## Abstract

Interferometric scattering (iSCAT) microscopy enables the label-free observation of biomolecule interactions with surfaces. Consequently, single-particle imaging and tracking with iSCAT is a growing area of study. However, establishing reliable cover glass passivation and functionalization methods are crucial to reduce non-specific binding and prepare surfaces for *in vitro* single-molecule binding experiments. Existing protocols for fluorescence microscopy can contain strongly scattering or mobile components, which make them impractical for iSCAT-based microscopy. In this study, we characterize several different surface coatings using iSCAT. We present approaches for cover glass passivation using 3-aminopropyltriethoxysilane (APTES) and polyethylene glycol (PEG, 2k) along with functionalization via a maleimide-thiol linker. These coatings are compatible with water or salt buffers, and show low background scattering; thus, we are able to measure proteins as small as 60 kDa. In this technical note, we offer a surface preparation suitable for *in vitro* experiments with iSCAT.

## Introduction

Interferometric scattering (iSCAT) microscopy is a label-free technique relying on common-path interferometry between the light scattered by single molecules and light reflected by the coverslip. Imperfections in the coverslip produce a high and spatially non-uniform background in iSCAT measurements. Strongly scattering objects such as gold nanoparticles offer a strong enough signal to be directly distinguished from the background^1^. In contrast, the signal of contrast, the signal of weakly scattering objects such as single proteins are indistinguishable from the background. Taking advantage of the temporally static background, ‘moving average’ background subtraction methods, also referred to as temporal filtering, enable signals from single proteins and small protein complexes to be detected in the process of adsorbing onto, desorbing from, or moving along a coverslip^2–6^. With single-molecule sensitivity, iSCAT has proven valuable for measuring the molecular mass and oligomerization states of single biomolecules^2^ via mass photometry and for detecting the dynamics of biomolecules such as motor proteins^7^, protein filaments^8^, DNA on two-dimensional materials^9^ and viruses and proteins on supported lipid bilayers (SLB)^3–5^.

In single-molecule fluorescence studies, strategies have been established to engineer the surface for investigating protein-protein and protein-nucleic acid interactions. Because non-specific adsorption introduces noise, surface passivation is often used to reduce direct binding to the surface. Effective strategies include blocking with non-fluorescent proteins^10^ or coating the surface with polyethylene glycol (PEG)^11,12^. Once the surface is passivated, immobilization of a target molecule onto the surface is used to observe repeated and specific binding to the same molecule or protein assembly^13^. Surface functionalization methods often employ the strong non-covalent avidin-biotin interaction to adhere target proteins or nucleic acid to the surface^14–16^.

It would be of great interest to perform single-molecule binding assays with iSCAT, to study the assembly of proteins on cytoskeletal elements, or on DNA. However, most of these passivation and functionalization approaches are not directly applicable to iSCAT. Without careful design, the signal from coatings can be difficult to distinguish from that of the molecules of interest, because proteins and long polymers can introduce a dynamic scattering background. For example, proteins or polymers blocking the surface could detach under strong iSCAT illumination conditions, and long PEG brushes have intrinsic mobility^17,18^ and strong scattering with a signal that scales with the molecular weight (MW)^2,6^. In contrast, the ideal surface passivation for iSCAT microscopy would be temporally static, and inert to light and non-specific interactions, offering a sufficiently high signal-to-noise ratio (SNR) for single biological molecules. Additionally, the ideal surface functionalization allows the formation of an adequately strong bond to anchor a target biomolecule under conditions (e.g. pH, temperature) that maintain the biological activity of the protein or nucleic acid. To date, few studies have addressed the issue of surface passivation and functionalization for iSCAT microscopy^19^.

Here, we investigate several approaches for surface passivation and functionalization for iSCAT microscopy. We tested the stability and effectiveness of passivation methods with PEG and APTES after incubation with different commonly used buffers. We also implemented a surface preparation which presents maleimide-thiol functionalization for anchoring specific molecules. The coating offers a well-passivated surface with low background in iSCAT microscopy, making it a promising preparation for future single-molecule studies.

## Results

### Surface passivation for iSCAT measurements

#### PEG 30k

One of the most common methods for preventing non-specific binding is coating the surface with the polymer PEG. For *in vitro* fluorescence studies, PEG of MW 5k-8k g/mol was used when studying single-molecule dynamics of chromatin^20^ or super-resolving DNA origami structures^21^, while PEG of higher MW (30k) was used to improve the passivation quality when studying binding to reconstituted microtubules^14^.

We tested a surface coating with silane-PEG 30k. This is a long PEG polymer functionalized with a silane group that can form a covalent bond with the exposed hydroxyl groups of the coverslip (Figure 1A, SI). When measured with iSCAT, we observed a strong dynamic signal in deionized water or a high-salt buffer (150 mM NaCl in HEPES buffer) (Figure 1B). Interestingly, when analyzed for scattering contrast as is typically done in mass photometry^2^, the sample incubated with deionized (DI) water revealed nearly equal numbers of binding and unbinding events, indicated by events in the histogram with negative and positive contrast (Figure 1C). In the case of incubation with the high-salt buffer, we mainly observed unbinding events, corresponding to positive contrast, and of a lower contrast than in DI water. Thus, this surface did not appear promising for single molecule iSCAT.

**Figure 1:**
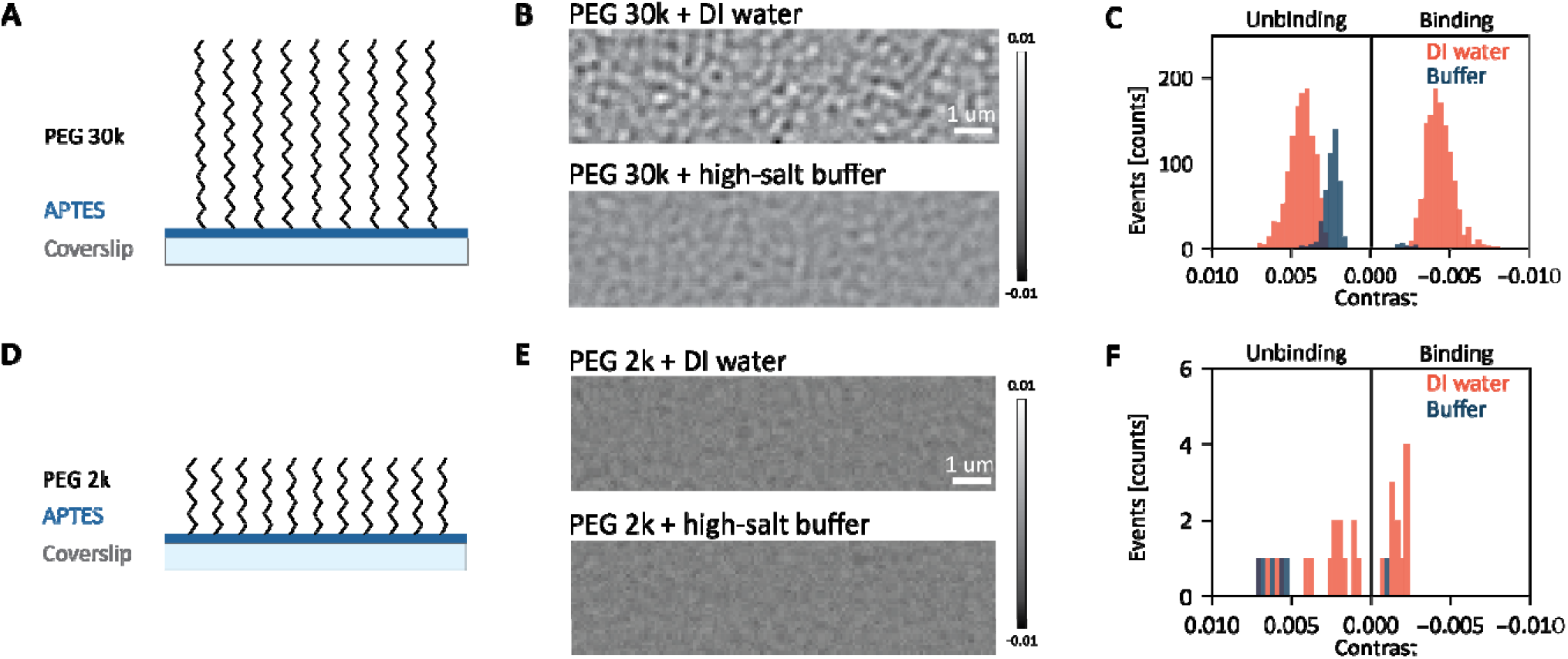
Passivation approaches for iSCAT measurements. (A) Sketch of coverslip coated with APTES and PEG 30k. (B) Examples of ratiometric iSCAT data of coverslips coated with PEG 30k and incubated with different buffers. Scale bar is 1 μm. (C) Example histogram of adsorbing (N(DI water)=1230, N(high-salt buffer)=469) and desorbing (N(DI water)=1226, N(high-salt buffer)=28) events detected on data shown in (B), 6000 frames. (D) Sketch of coverslip coated with APTES and PEG 2k. (E) Examples of ratiometric iSCAT dat of coverslips coated with PEG 2k and incubated with different buffers. Scale bar is 1 μm. (F) Histogram of adsorbing and desorbing events detected on data shown in (E). Data includes 3 measurements on 3 different coverslips with each 3000 frames.

#### PEG 2k

Aiming to decrease the signal of the passivated surface, we tested PEG of a lower MW (2k). We envisaged that this would lead to less scattering, and also reduced mobility of the PEG brushes due their shorter length. For experiments with PEG 2k, we used a 3-aminopropyltriethoxysilane (APTES) surface coating and its reaction with succinimidyl ester-functionalized PEG^12^. Surfaces with PEG 2k demonstrate reduced background signal and a low number of detected events (Figure 1D-F) compared to PEG 30k. The surface shows no evidence of unbinding events from PEG detachment from the coverslip, even when incubated with high-salt buffer.

To confirm the presence of PEG on the surface, we employed fluorescently labeled PEG-Cy3 (Figure S2). We carried out a titration, varying the weight percentage of PEG-Cy3 (0%, 5%, 10%, and 15%). The detected fluorescence intensity appears nearly uniform across the field of view, and increases linearly with the weight percentage of fluorescent PEG, as expected. Additional photobleaching experiments demonstrated the stability of PEG surface layer during an eight-minute DI water incubation (Figure S6). The liquid deposition of APTES led to less uniform, but higher surface coverage compared to vapor-phase deposition (Figure S8).

#### Mass photometry on a passivated surface

To measure the scattering signal from proteins relative to the PEGylated (2k) surface, and to test the passivation quality, we incubated the surface with different proteins (Figure 2A,B): neutravidin (60 kDa), IgG (160 kDa), and fibrinogen (340 kDa, dimer). Individual binding events are clearly visible in the temporally filtered data(Figure 2C).

**Figure 2:**
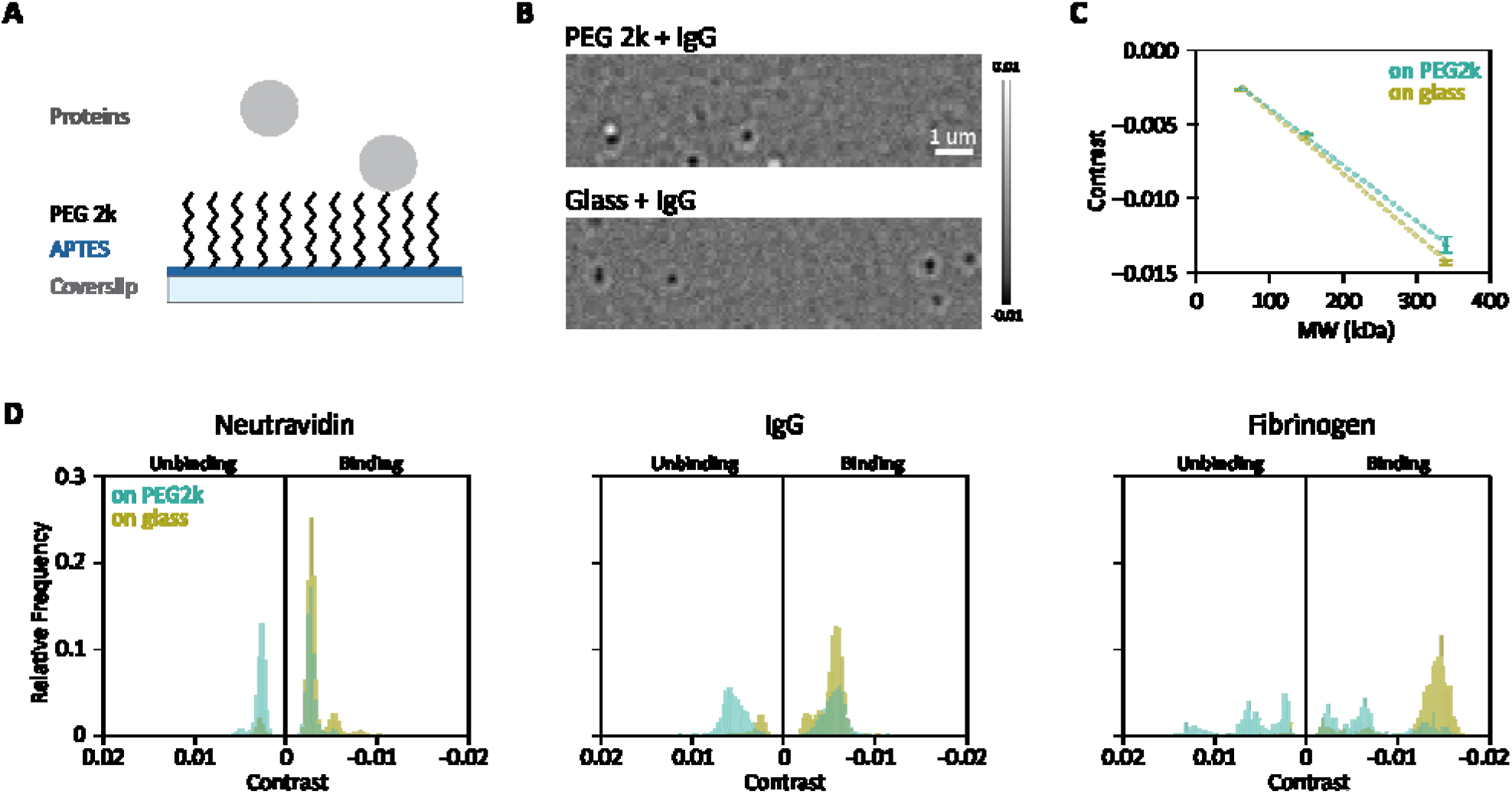
Mass photometry on passivated surface. (A) Sketch of coverslip coated with APTES and PEG 2k incubated with proteins. (B) Examples of ratiometric iSCAT data of coverslips coated with PEG 2k or just glass and incubated in DI water with 10 nM IgG. Scale bar is 1 μm. (C) Mass calibration on PEG 2 kDa and glass surfaces using Neutravidin monomer, IgG and Fibrinogen (60, 150 and 340 kDa respectfully). (D) Histograms of events detected on glass and PEG 2kDa surfaces with neutravidin (10 nM on glass, 15 nM on PEG), N(PEG, binding events)= 2197, N(PEG, unbinding events)= 1700, N(glass, binding events)=4670; IgG (10 nM on glass, 15 nM on PEG), N(PEG, binding events)= 3780, N(PEG, unbinding events)= 3682, N(glass, binding events)= 4322; Fibrinogen (10 nM on glass, 30 nM on PEG), N(PEG, binding events)= 322, N(PEG, unbinding events)= 166, N(glass, binding events)= 565. For neutravidin and IgG, data shows three merged measurements per condition (6000 frames each) as 1 experimental replicate. For fibrinogen, data shows five merged measurements per condition (12000 frames each).Two more replicates are highlighted in S9. Relative frequency refers to the number of events normalized to the total number of events per condition.

The histograms of ratiometric contrast for neutravidin, IgG and fibrinogen appear nearly symmetrical on a PEGylated surface, implying equal numbers of detected binding and unbinding events and confirming a high-quality passivation (Figure 2C). As a control, we measured the proteins on untreated glass. As expected, the glass surface exhibits exclusively binding events, since molecules remain at the surface rather than unbinding. These differences between PEG 2k and bare glass surfaces are also apparent when we plot the number of binding and unbinding events per frame (Figure S5). While PEG 2k reveals nearly equal numbers of binding and unbinding events per frame for each protein, leading to a binding/unbinding ratio between 1 and 2 for the entire 1-2-minute measurement (Table S4), proteins bind to the glass surface with a decaying number of events per frame and a very low number of unbinding events. This emphasizes that typical mass photometry measurements on glass are performed in a manner that is out-of-equilibrium, so that the rate of binding changes with time as proteins fall out of solution and stick irreversibly to the coverslip. Nevertheless, both surfaces yield similar mass calibration curves (Figure 2D).

Surface passivation, as detailed above, plays a pivotal role in minimizing nonspecific binding to the coverslip. Having established this foundation, in the subsequent section we delve into methods aimed at surface functionalization, to allow target molecules to be displayed on the surface.

### Surface functionalization for iSCAT measurements

#### Poly-L-lysine (PLL)

In mass photometry, molecules must dwell at the surface to produce a scattering signal and be measured. Due to the negative charge of bare glass surfaces, negatively charged biomolecules such as DNA require surface preparation. Homopolypeptide coatings, such as PLL, among others, have been used to create positively charged surfaces^22,23^. Experiments are typically carried out over short time periods (up to 60 s) and with low-salt buffers.

We investigated the use of PLL as a coating to allow long-term DNA adherence (Figure 3A). To test the long-term stability of the PLL coating under different buffer conditions, we prepared PLL-coated coverslips. We incubated the surface with either DI water or high-salt buffer, imaged the sample at different time points (0, 5, 10 and 20 min) after initial incubation and counted the detected events (see SI). While the number of events for PLL coated slides incubated with DI water remained negligible over the measured time range, significant activity appeared when slides were incubated with high-salt buffer (Figure 3B,C). It is typical to detect some events (< 100 / 60 sec), even in measurements with only a clean coverslip and buffer (Figure S9), but this is more than an order of magnitude greater than those rates. We speculate that the high ionic strength buffer leads to a disintegration of the coating, which appears as particles in solution. Thus, PLL coatings are stable under iSCAT illumination conditions with DI water, even over long-time ranges, but they are not suitable in combination with high-salt buffers.

**Figure 3:**
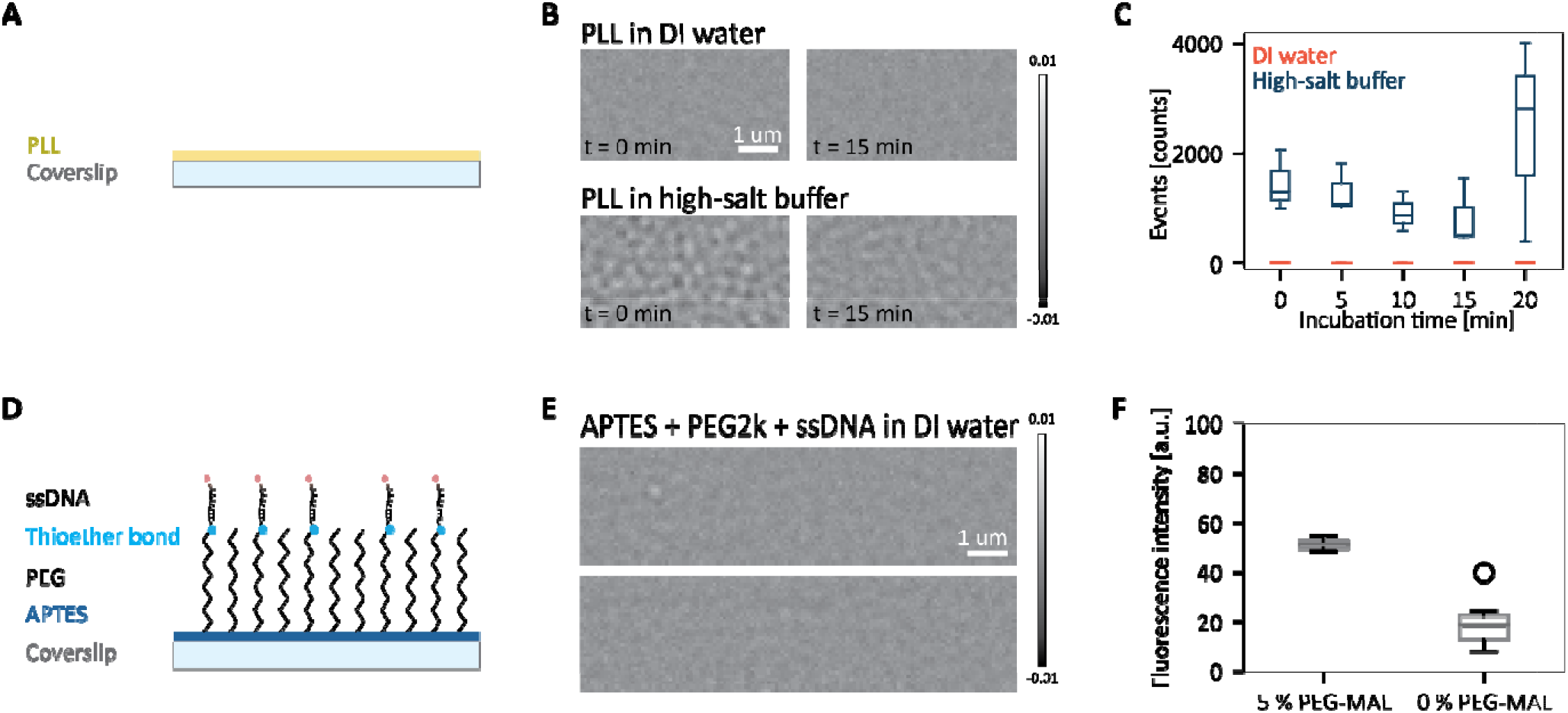
Functionalization approaches for iSCAT measurements. (A) Sketch of PLL-coated coverslip. (B) Examples of ratiometric iSCAT data of PLL-coated coverslips incubated with different buffers. (C) Number of total events (adsorbing and desorbing) detected. Number of samples for each condition: N = 3; 6000 frames each. (D) Sketch of a coverslip coated with APTES, SVA-PEG-Mal, Thiol-DNA-Atto633. (E) Examples of ratiometric iSCAT data of PEG2k coated coverslips functionalized with SH-DNA-Atto633. (F) Mean fluorescence intensity of FOV of coverslips coated with 5 % PEG-Mal and 95 % PEG or 0 % PEG-Mal and 100 % PEG after incubation with SH-DNA-Atto633, N = 6.

#### PEG-Maleimide and Thiol-DNA

Another approach using covalent bonds to functionalize surfaces employs thiol-maleimide chemistry to form a thioether bond. Since linkages between maleimide and thiol molecules are stable at physiological pH conditions, this method is commonly used for protein labelling as well as protein cross-linking via the thiol group contained in cysteine residues of proteins^24,25^.

In our case, we used maleimide-functionalized PEG (PEG-Mal) and DNA oligonucleotides modified with a thiol group (thiol-DNA) (Figure 3D). In iSCAT measurements of these fully functionalized surfaces, we observe a signal comparable to that for PEG2k-coated surfaces (Figure 3E, S11). To validate the presence and stability of the thioether bond between PEG and DNA, we used thiol-DNA labelled with a fluorescent dye (Atto633). We changed the concentration of PEG-Mal (0 and 5 % PEG-Mal), while keeping the concentration of thiol-DNA constant (Figure 3F). The fluorescence intensity of the PEG-Mal functionalized surface increased compared to the control group lacking functionalized PEG, providing evidence of DNA binding via thioether bonding. Additional photobleaching experiments demonstrated the stability of the thioether bond during an eight-minute DI water incubation (Figure S10).

## Discussion

Surface passivation with PEG can be compatible with iSCAT measurements. We found that smaller molecular weight PEG produced a smaller dynamic background signal in iSCAT measurements.

PEG30k shows a strong dynamic signal dependent on the buffer solution. We suspect that in DI water, due to the lower ionic strength, the PEG brush conformation allows for more flexibility – whereas in high ionic strength conditions, they are more densely packed^26^. In high-salt buffer, we mainly observe molecules leaving the surface – perhaps of smaller fragments released by thermal-catalyzed hydrolysis which PEG is known to suffer from^27,28^. In contrast, PEG2k measurements have a dynamic background signal comparable to glass measurements. This could be due to improved attachment, or due to the shorter chain length which if fragmented would not scatter much. Overall, large molecular weight PEG scatters too much to be used in studies of single proteins but could be useful when studying strongly scattering objects such as gold nanoparticles. With PEG 2k, we demonstrated measurements of proteins down to 60 kDa. Even smaller PEG might further decrease the mass limit, but would require validation that the required passivation quality is achieved.

In mass photometry measurements on PEG2k-passivated surfaces we noticed a greater variance in contrast values compared to untreated glass for IgG, but less so for fibrinogen and neutravidin. We quantified these differences by fitting a Gaussian to the main peak in the contrast histogram (Table S4). We hypothesize that this difference in variance is a result of contributions from scattering from the PEG and proteins moving along the surface before they bind (motion blur). Proteins which only bind or unbind during the temporal filtering window should appear as a black or white signal respectively. As we report, IgG proteins interacting with the PEG 2k surface bind and unbind at equal rates, and some proteins move laterally while they are in contact with the surface, creating a black spot with a white tail in the temporally filtered images. The black and white signals are analyzed as separate binding and unbinding events with more variable contrast than the stationary protein. Further development on the effect of the temporal filtering methods could be used to correct this effect.

We also demonstrated that the PEG layer used for passivation can be functionalized at physiological pH and room temperature, allowing anchoring of biological molecules. Using PEG-Maleimide creates a surface with an available thiol-group, allowing the potential to bind various biological molecules. We tested the surface with thiol-DNA, and found it to offer a similar low-scattering background to PEG 2k.

Here, we used a sensitive setup intended for mass photometry to characterize passivated and functionalized surfaces. We thus expect that these surfaces will be compatible with less sensitive iSCAT setups. We envision that this study will not only help future mass photometry and iSCAT microscopy studies, but can also be used in other label-free microscopy techniques.

## Acknowledgements

We thank Maartje Bastings and Beat Fierz for helpful discussions. We acknowledge Kaltrina Paloja, Jorieke Weiden and Alice Comberlato for generously providing us samples of DNA origami and buffers and for useful discussions. We thank Charlotte Aumeier for the PEGylation protocol and training. We thank Kyle Douglass for technical editing and proofreading. This work was supported by the National Centre for Competence in Research (NCCR) Chemical Biology.

## Supporting Information

- PDF with Materials and Methods and additional experimental results
- Movie 1: Silane-PEG 30k-coated coverslips incubated with DI water and high-salt buffer
- Movie 2: PEG2k-coated coverslips with DI water and 10 nM IgG

## Author contribution

J.S. and L.E. conducted experiments, analyzed the data, and prepared the figures. J.S., L.E. and S.M designed the experiments and wrote the manuscript.

## Notes

### Competing Interest Statement

The authors have declared no competing interest.

### Summary of Updates

Figure 2 revised; Supplemental files updated

