## Supplementary Information for "Surface passivation and functionalization for interferometric scattering microscopy"

### Materials and Methods

#### Materials

Coverslips (24 x 50 mm, Marienfeld, 0101222), NAP-10 size exclusion chromatography column (NAP™-10 20 ST, Cytiva, GE17-0854-01), silicone gaskets (3 mm × 1 mm, Grace Bio-Labs, GBL103250).

PLL solution (0.01 %, Sigma-Aldrich, P4832), (3-Aminopropyl)triethoxysilane (APTES 99 %, Sigma-Aldrich, 440140), Methanol (≥99%, Fischer Scientific GmBH, 10511732)acetic acid (100% (glacial), Merck kGaA, 100063), DL-Dithiothreitol (DTT, Sigma-Aldrich, D9779), Glutaraldehyde (GA, 25 % in $H_2O$, Sigma-Aldrich, G5882), Magnesium chloride solution (1 M, Sigma-Aldrich, M1028), HEPES buffer (1 M, Chemie Brunswig, FIBBP299-500), Sodium chloride solution (5 M, Sigma-Aldrich, S5150), Sodium bicarbonate (Sigma-Aldrich, S5761), IgG (Jackson ImmunoResearch, 712-005-150), Neutravidin (Thermo Scientific 10443985), Fibrinogen (Sigma-Aldrich, F3879).

mPEG-SVA (2 kDa, LaysanBio, MPEG-SVA-2000-1g), SVA-PEG-Mal (3.4 kDa, LaysanBio, MAL-PEG-SVA-3400-1gram), Cy3-PEG-NHS (2 kDa, Nanocs, PG2-NSS3-2k), tri-ethoxy-silane-PEG (30 kDa, Creative PEGWorks, PBS-2014), mPEG-NHS (333 Da, VWR, TCM2186-25MG).

DNA sequences were purchased from Integrated DNA Technologies (IDT): DNA Dithiol (sequence:TTATACATCTA/3'Dithiol), SH-DNA-Atto633 (sequence: 5'Atto633/TTATACATCTA/3'Dithiol), Amine-DNA-Cy5 (sequence: /5AmMC6/CT AGA TGT AT/3Cy5Sp/), Amine-DNA (sequence: /5AmMC6/CT AGA TGT AT)

DNA origami, previously described by Eklund et al.[1], was modified with six amine handles on one side and labelled with six Cy5 dye molecules. Another version was only modified with the dye, no amine handles were added.

#### Buffer preparation

The following buffers were used for sample preparation and imaging: $MgCl_2$-enriched HEPES buffer (high-salt buffer): 10 mM HEPES, 150 mM NaCl, pH7.4, 18 mM $MgCl_2$. Sodium bicarbonate buffer: 100mM, pH 8.5.

#### Coverslip cleaning

Coverslips were cleaned according to the protocol outlined in the Refeyn Manual for OneMP. Briefly, the coverslips were cleaned by sequentially placing in an ultrasound bath with Hellmanex (2 %), DI water, isopropanol and DI water and drying using a nitrogen stream. Between each sonication step three washing steps with DI water were performed.

#### PLL coating

We followed a protocol provided by Refeyn[2]. Briefly, 7 µl of PLL solution was sandwiched between two coverslips. After 30 seconds, the coverslips were extensively washed with DI water and subsequently dried with nitrogen. The functionalized coverslips were used within the same day.

#### APTES + Glutaraldehyde (GA) coating and functionalization with amine-DNA

Coverslips were vapor-deposited with the aminosilane APTES (3h, described below). Subsequently we assembled a flow chamber (described below) and incubated with 5 % Glutaraldehyde (GA) in water for 2 hours. For the titration experiment, after washing the samples were incubated with 6x amine-functionalized DNA origami[1] (0, 1 or 13 nM in high-salt buffer, overnight) and subsequently washed. Imaging was executed on the same day.

#### Silane-PEG coating

We used a protocol described in Aumeier et al.[3].

#### APTES + PEG coating and functionalization with thiol-DNA

The PEGylation protocol for coverslips was adapted from Chandradoss et al.[4] Briefly, we activated the surface using oxygen plasma (Diener Pico, 160 W, 300 s).

For liquid APTES deposition, the coverslips were immediately immersed in methanol. Subsequently, coverslips were immersed in a solution of methanol (>=100%, 100 mL), APTES (3 mL), and acetic acid (5 mL), rinsed with methanol, and were then dried using a nitrogen stream.

Alternatively, for gas APTES deposition, a few drops (3- 5) of APTES were dispensed into a petri dish, and coverslips were positioned in a holder above, within a vacuum desiccator. The coverslips were left under vacuum for 5 minutes before being removed.

Following silanization, we used mixtures of PEG 2k for PEGylation. In case of experiments with PEG-Cy3, we mixed NHS-PEG-Cy3 with mPEG-SVA (10% by weight, unless otherwise stated). In case of experiments with Maleimide, mPEG-SVA was doped with SVA-PEG-Mal (5 % by weight). For each coverslip, 8.2 mg of the solid PEG mixture was dissolved in freshly prepared sodium bicarbonate buffer (64 µl, pH 8.5), followed by gentle pipetting up and down. The PEG solution underwent centrifugation (16 100 x g, 1 min). Subsequently, coverslips were placed in a humid chamber, and the prepared PEG solution (70 µl) was added. Another silanized slide was then added on top, creating a coverslip 'sandwich'. The coverslips were left to incubate overnight in a humid chamber. Subsequently, coverslips were thoroughly washed with DI water and dried using nitrogen.

For experiments using double PEGylation, the protocol described in Chandradoss et al. was followed[4].

We noticed two critical points: (1) When assembling the coverslip 'sandwich', ensure the top coverslip is lowered gently, taking care to avoid the formation of air bubbles. (2) For humid chamber preparation, saturate a piece of paper towel with DI water, and place it into a box with a lid. Place a sheet of parafilm on top of the towel, leaving some space uncovered to allow water to evaporate inside the box. Arrange coverslips on the parafilm layer and prepare them into the 'sandwich' as described above. Close the lid and leave to incubate.

At this point, the coverslips could be used as an APTES-PEG surface, or further functionalized.

To conjugate DNA with the surface via thiol-maleimide chemistry, we followed these steps: Because the thiol group would be unstable over time, the DNA oligonucleotides were purchased modified with a dithiol group, which we subsequently reduced to the active thiol group immediately prior to usage according to a Sigma-Aldrich protocol[5]. In brief, the dithiol-functionalized DNA oligonucleotide was incubated in a solution of DTT (100 mM) for 1 hour. The resulting reduced product was then collected using a NAP-10 size exclusion chromatography column.

The concentration of the newly formed DNA-thiol solution was measured using NanoDrop 2000 (Thermo Scientific). The solution was then diluted with deionized water (DI water) to a concentration of 1.82 µM. This diluted solution (64 µl) was added to the previously dried PEGylated coverslips, forming the previously described coverslip 'sandwich', and left to incubate in a humid chamber at 4 °C overnight. The following day, samples were washed with DI water, dried with a nitrogen stream and imaged on the same day.

#### Coverslip preparation

Different experiments utilized either flow chambers or gaskets. Gaskets should only be used for short measurements, as evaporation may occur if imaged for an extended period.

Flow chambers: Flow chambers were prepared by sticking double-sided tape between a functionalized coverslip and a cleaned coverslip. The flow chamber was initially filled with buffer. The specimen was loaded using a filter paper. After adjusting the focus of the microscope, the measurement was started. Flow chambers were used in all fluorescence experiments and iSCAT experiments with the following coatings: Silane-PEG-30k and APTES+GA.

Gaskets: The silicone gaskets were attached to the top of the coverslip (4x4 array). The buffer solution was added before imaging. After adjusting the focus of the microscope, the sample solution was added to the gasket, mixed by pipetting up and down and then directly imaged (typical lag time around 30 s). Gaskets were used in iSCAT experiments with 2 kDa PEG or PEG-Mal surface and with bare glass.

#### Protein sample preparation

All proteins used were diluted to 150 nM in HEPES buffer (20 mM) before imaging. During coverslip preparation, DI water was used and the prepared protein sample was added as described in the section above (coverslip preparation). The concentration of protein used for imaging on glass was 10 nM (18.67 µl and 1.33 µl of DI water and protein respectively). For imaging proteins on PEG, final concentrations used were: IgG (10 nM on PEG and glass), Neutravidin (10 nM on glass, 15 nM on PEG) and Fibrinogen (10 nM on glass, 30 nM on PEG). The volumes of buffer and protein sample we added in each case were dependent on the required final concentration, ensuring a final volume of 20 µl.

##### iSCAT imaging

iSCAT imaging was performed on the Refeyn One MP microscope, with data acquisition conducted using the Acquire MP (AMP) software. A field of view of 128 x 34 pixels (pixel size = 84.4 nm) was imaged with a frame rate of 994 Hz, effectively becoming 99.4 Hz after temporal binning. Measurements included 6,000 frames (12,000 for fibrinogen on PEG2k measurements) unless otherwise stated.

Data analysis was carried out using the Refeyn Discover MP (DMP) software. Parameters used were T1 = 1.500, T2 = 0.250, W = 2.3085750000, A12 = -5.3989470000 and S = 5.2816720000. Extracted events were further processed using Python.

##### Fluorescence imaging

Fluorescence data was acquired using a fluorescence widefield microscope (Zeiss) equipped with a 63x oil immersion objective (NA 1.4) and a sCMOS camera (PRIME BSI, Photometrics). Prior to all fluorescence imaging, flow chambers were assembled following the procedure described in the previous section. All experiments were conducted with an exposure time of 100 ms.

For photobleaching experiments, a FRAP laser (405nm, 25 mW, 2D-VisiFRAP, Visitron Systems GmBH) was used to photobleach a small area within the field of view. Subsequently, 500 frames were acquired with a time interval of 1 s between each frame. Mean and median fluorescence intensities of the photobleached area were analyzed using ImageJ[6].

Fluorescence measurements of PEG-Cy3 were imaged with excitation by a 550 nm LED (pE-800 LED Illumination System, CoolLED) at 30 mW power, those of SH-DNA-Atto633 were imaged with a 635nm, 30 mW LED light source (pE-800 LED Illumination System, CoolLED). In both cases a multi-band pass filter (Chroma 89402) was utilized to separate the illumination and emission.

##### Data analysis

Analysis of PEG-Cy3 titration experiment: ImageJ software was used to analyze mean fluorescence values from generated data. A region of interest (ROI) was cropped out in all images (1000 x 1000 pixels) in the center of the imaged field of view.

Photobleaching experiments analysis: The photobleached area was selected as a ROI on ImageJ. Mean and median fluorescence intensity data was derived from values within the photobleached ROI before and after photobleaching.

#### Supplementary Notes

##### S1. iSCAT measurements of PEG30k-coated surface

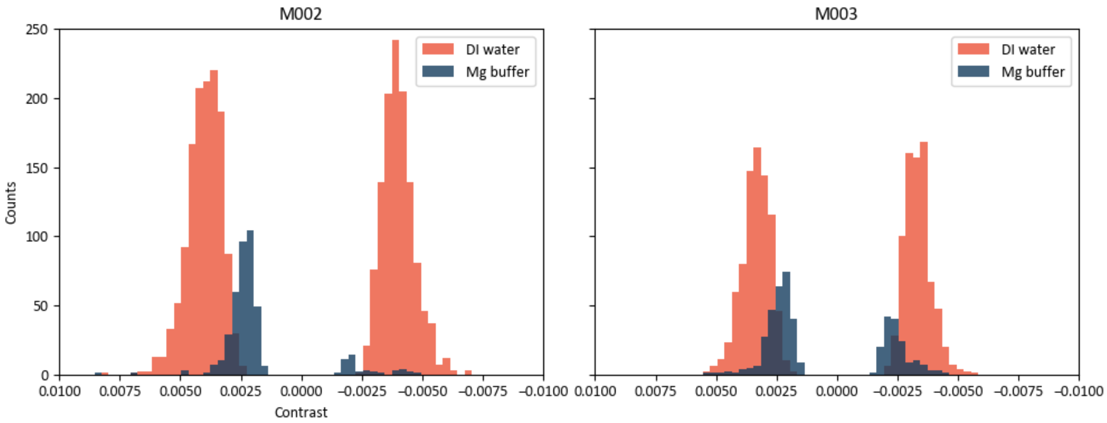

**Figure S1.** iSCAT measurements of PEG30k-coated surface. This figure shows two replicate measurements of the experiment illustrated in Figure 1C. M002: histogram of adsorbing (N(DI water)=1215, N(high-salt buffer)=44) and desorbing (N(DI water)=1329, N(high-salt buffer)=367) events. And M003: histogram of adsorbing (N(DI water)=769, N(high-salt buffer)=160) and desorbing (N(DI water)=811, N(high-salt buffer)=282) events. Note that the high-salt buffer is named “Mg buffer” in the figure.

#### S2. Titration of PEG-Cy3.

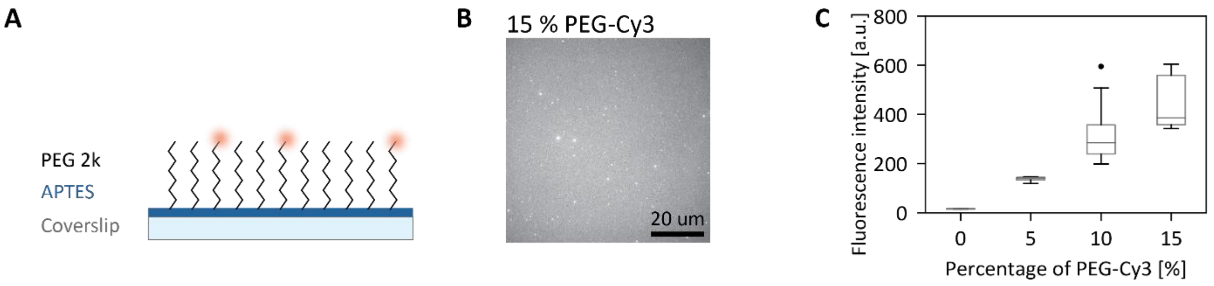

**Figure S2.** Titration of PEG-Cy3. (A) Illustration of surface coated with APTES and PEG2k/PEG2k-Cy3 mixture. (B) Example fluorescence image. (C) Mean fluorescence intensity per FOV of samples with different PEG-Cy3 fractions. Number of FOV imaged on 2 coverslips per condition (0%, 5%, 10%, 15% PEG-Cy3) = (6, 8, 16, 7).

#### S3. PEG 2kDa passivation with 3 proteins

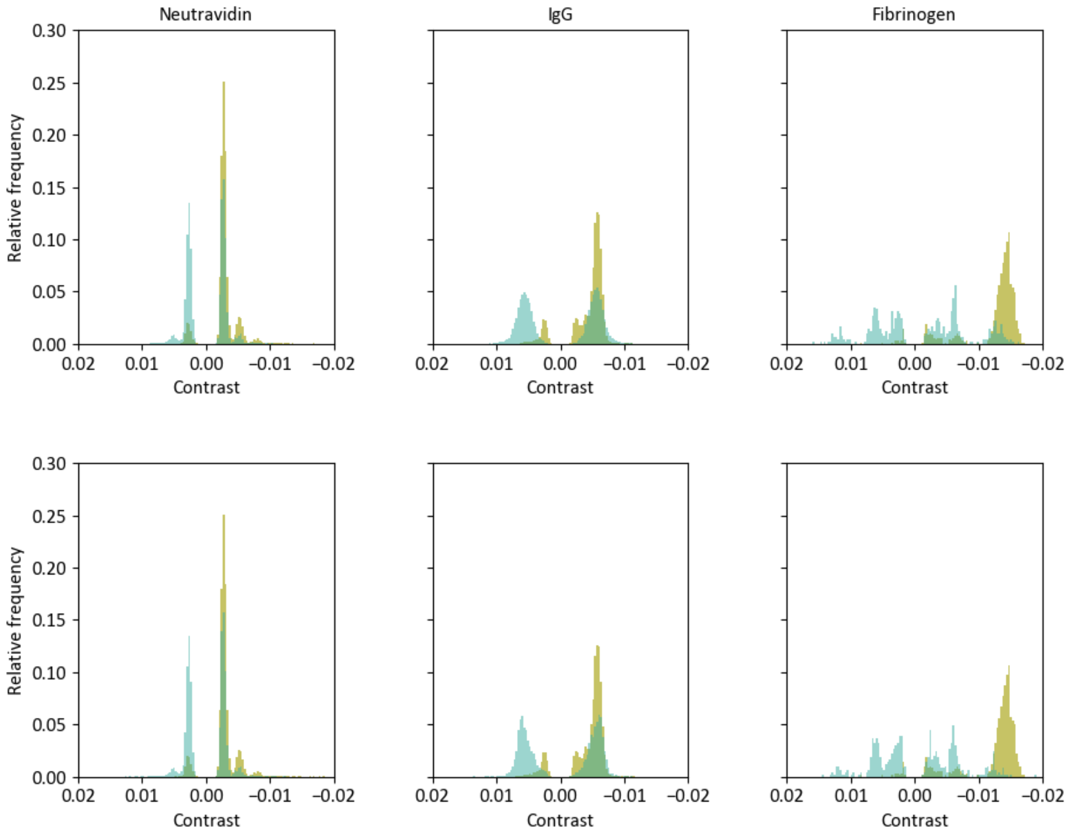

**Figure S3.** Histograms of events detected on glass and PEG 2kDa surfaces using 3 proteins: Neutravidin, IgG and Fibrinogen. This figure shows two additional experimental replicates for data shown in Figure 2C in the main text, with 3 measurements per condition. In yellow is the control measurement on glass. In green the measurements on PEG 2k. N represents counts under the Gaussian. Row 1: N (Neutravidin glass binding) = 5377, N (Neutravidin PEG binding) = 1371, N (Neutravidin PEG unbinding) = 1094, N (IgG glass binding) = 4322, N (IgG PEG binding) = 7855, N (IgG PEG unbinding) = 7562, N (Fibrinogen glass binding) = 565, N (Fibrinogen PEG binding) = 265, N (Fibrinogen PEG unbinding) = 161. Row 2: N (Neutravidin glass binding) = 5377, N (Neutravidin PEG binding) = 2035, N (Neutravidin PEG unbinding) = 1761, N (IgG glass binding) = 4322, N (IgG PEG binding) = 3044, N (IgG PEG unbinding) = 2935, N (Fibrinogen glass binding) = 565, N (Fibrinogen PEG binding) = 125, N (Fibrinogen PEG unbinding) = 89.

#### S4. Analysis from mass photometry measurements

| | Ratiometric Contrast | | | | | | Ratio (binding/unbinding) |
|---|---|---|---|---|---|---|---|
| | *Binding* | | | *Unbinding* | | | |
| **Measurement** | **Mean** | **Sigma** | **Counts under the gaussian** | **Mean** | **Sigma** | **Counts under the gaussian** | |
| IgG on glass | -0.00578 | 0.0006 | 4322 | | | | |
| IgG on PEG replicate 1 | -0.00558 | 0.0014 | 3780 | 0.00553 | 0.00138 | 3682 | 1.0266 |
| IgG on PEG replicate 2 | -0.00562 | 0.0014 | 7855 | 0.00562 | 0.00140 | 7562 | 1.0387 |
| IgG on PEG replicate 3 | -0.00569 | 0.0012 | 3044 | 0.00568 | 0.00132 | 2935 | 1.0371 |
| Neutravidin monomer on glass | -0.00275 | 0.0004 | 4670 | | | | |
| Neutravidin dimer on glass | -0.00526 | 0.0005 | 707 | | | | |
| Neutravidin monomer on PEG replicate 1 | -0.00269 | 0.0004 | 2197 | 0.00271 | 0.00039 | 1700 | 1.2924 |
| Neutravidin dimer on PEG replicate 1 | -0.00500 | 0.0006 | 114 | 0.00480 | 0.00061 | 129 | 0.8837 |
| Neutravidin monomer on PEG replicate 2 | -0.00268 | 0.0004 | 1264 | 0.00264 | 0.00035 | 994 | 1.2716 |
| Neutravidin dimer on PEG replicate 2 | -0.00512 | 0.0005 | 107 | 0.00513 | 0.00063 | 100 | 1.0700 |
| Neutravidin monomer on PEG replicate 3 | -0.00271 | 0.0003 | 1888 | 0.00269 | 0.00035 | 1592 | 1.1859 |
| Neutravidin dimer on PEG replicate 3 | -0.00509 | 0.0005 | 147 | 0.00490 | 0.00083 | 169 | 0.8698 |
| Fibrinogen on glass | -0.01425 | 0.001 | 565 | | | | |
| Fibrinogen on PEG replicate 1 | -0.01324 | 0.0013 | 322 | 0.01193 | 0.00115 | 166 | 1.9398 |
| Fibrinogen on PEG replicate 2 | -0.01276 | 0.0012 | 265 | 0.01178 | 0.00114 | 161 | 1.6460 |
| Fibrinogen on PEG replicate 3 | -0.01211 | 0.00150 | 125 | 0.01122 | 0.00139 | 89 | 1.4045 |

**Table S4-1:** Analysis of ratiometric contrast from measurements using PEG 2k surface on three proteins: Neutravidin, IgG and Fibrinogen.

| | Mass | | | | | | Ratio (binding/unbinding) |
|---|---|---|---|---|---|---|---|
| | *Binding* | | | *Unbinding* | | | |
| **Measurement** | **Mean** | **Sigma** | **Counts under the gaussian** | **Mean** | **Sigma** | **Counts under the gaussian** | |
| IgG on glass | 138 | 14.9 | 4322 | | | | |
| IgG on PEG replicate 1 | 140 | 36.0 | 3780 | -138 | 37.0 | 3682 | 1.0266 |
| IgG on PEG replicate 2 | 141 | 37.0 | 7855 | -141 | 37.0 | 7562 | 1.0387 |
| IgG on PEG replicate 3 | 143 | 33.0 | 3044 | -143 | 35.0 | 2935 | 1.0371 |
| Neutravidin monomer on glass | 64 | 8.7 | 4670 | | | | |
| Neutravidin dimer on glass | 125 | 12.9 | 707 | | | | |
| Neutravidin monomer on PEG replicate 1 | 63 | 10.1 | 2197 | -64 | 10.3 | 1700 | 1.2924 |
| Neutravidin dimer on PEG replicate 1 | 125 | 14.5 | 114 | -119 | 16.0 | 129 | 0.8837 |
| Neutravidin monomer on PEG replicate 2 | 63 | 9.4 | 1264 | -62 | 9.1 | 994 | 1.2716 |
| Neutravidin dimer on PEG replicate 2 | 128 | 12.4 | 107 | -130 | 16.8 | 100 | 1.0700 |
| Neutravidin monomer on PEG replicate 3 | 64 | 9.0 | 1888 | -63 | 9.2 | 1592 | 1.1859 |
| Neutravidin dimer on PEG replicate 3 | 127 | 13.8 | 147 | -122 | 22.0 | 169 | 0.8698 |
| Fibrinogen on glass | 343 | 25.0 | 565 | | | | |
| Fibrinogen on PEG replicate 1 | 342 | 34.0 | 322 | -308 | 30.0 | 166 | 1.9398 |
| Fibrinogen on PEG replicate 2 | 330 | 32.0 | 265 | -304 | 30.0 | 161 | 1.6460 |
| Fibrinogen on PEG replicate 3 | 312 | 40.0 | 125 | -289 | 37 | 89 | 1.4045 |

**Table S4-2:** Analysis of molecular mass from measurements using PEG 2k surface on three proteins: Neutravidin, IgG and Fibrinogen. Note that only the mean and sigma columns differ from Table S4-1, since the same datasets were analyzed and converted here to mass using the mass calibration curve.

### S5. Mass photometry: Binding and unbinding events per time

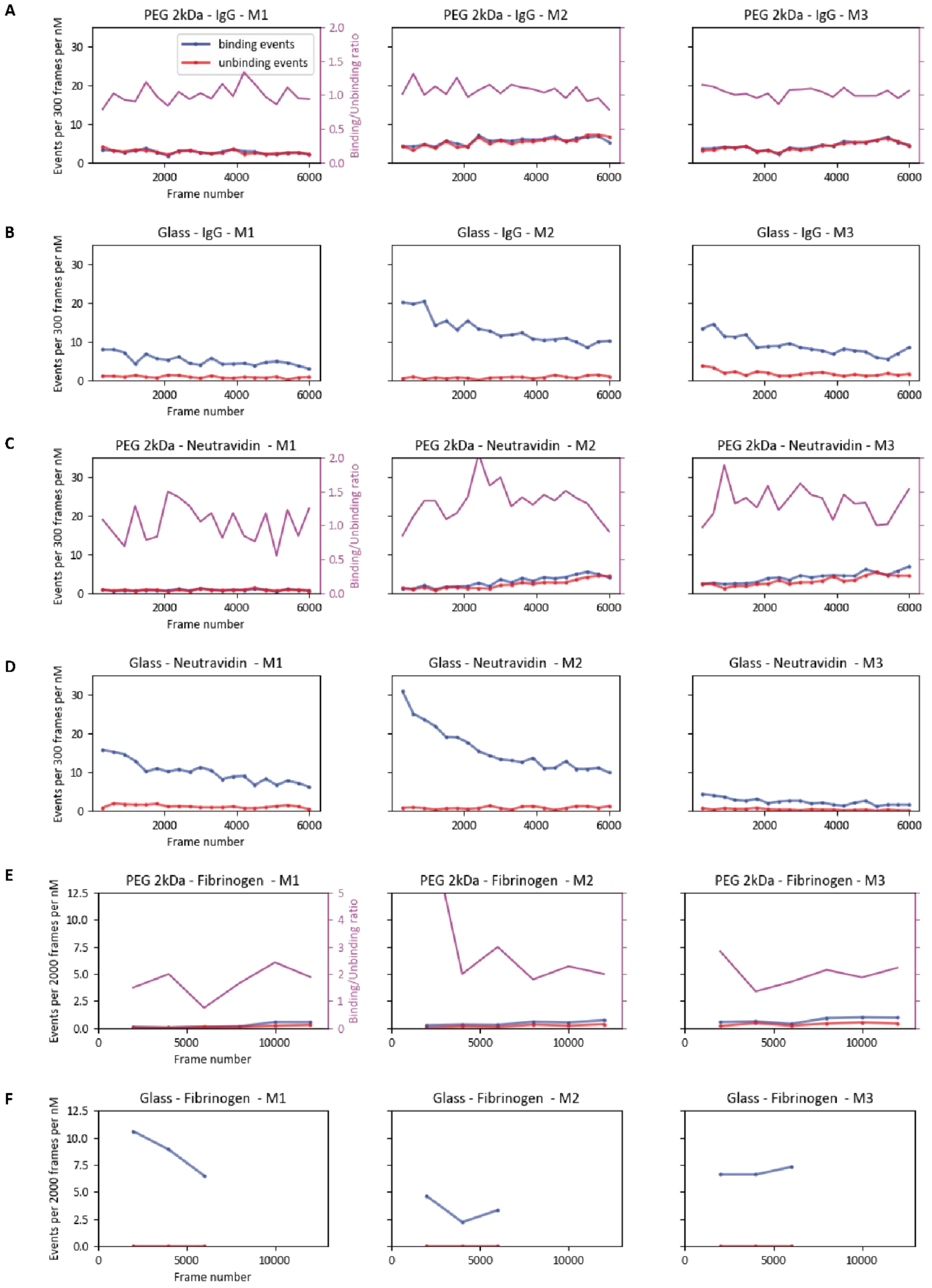

**Figure S5.** Binding and unbinding events per time, normalized by the protein solution concentration. (A) Three measurement of 15 nM IgG on PEG 2k. (B) Three measurements of 10 nM IgG on glass. (C) Three measurements of 15 nM Neutravidin on PEG 2k. (D) Three measurements of 15 nM Neutravidin on glass. (E) Three measurements of 30nM fibrinogen on PEG 2k (12 000 frames each). (F) Three measurements of 10nM fibrinogen on PEG 2k. For (E,F) we increased the bin width to 2000 to compensate for the reduced surface landing rates of fibrinogen.

#### S6. Photobleaching experiments for bond between APTES and NHS-PEG.

To test the stability of the bond formed between the APTES coating and the NHS-PEG, we prepared samples with 10 % fluorescent PEG 2k (PEG-Cy3, by weight) on top of coverslips with and without prior APTES coating. We compared the fluorescence intensity between silanized (coated with APTES and treated with NHS-PEG) and non-silanized (bare glass treated with NHS-PEG) surfaces and observed consistently a higher intensity for coverslips without APTES.

To further assess the stability of the formed bonds, recovery after photobleaching was measured for both the silanized and non-silanized PEG-Cy3 surfaces. The data reveals no recovery after photobleaching for 8 minutes (1 fps) in both silanized and non-silanized coverslips.

These results indicate that, under the experimental conditions used and independent of APTES coating, PEG 2k shows good stability within a duration of 8 minutes.

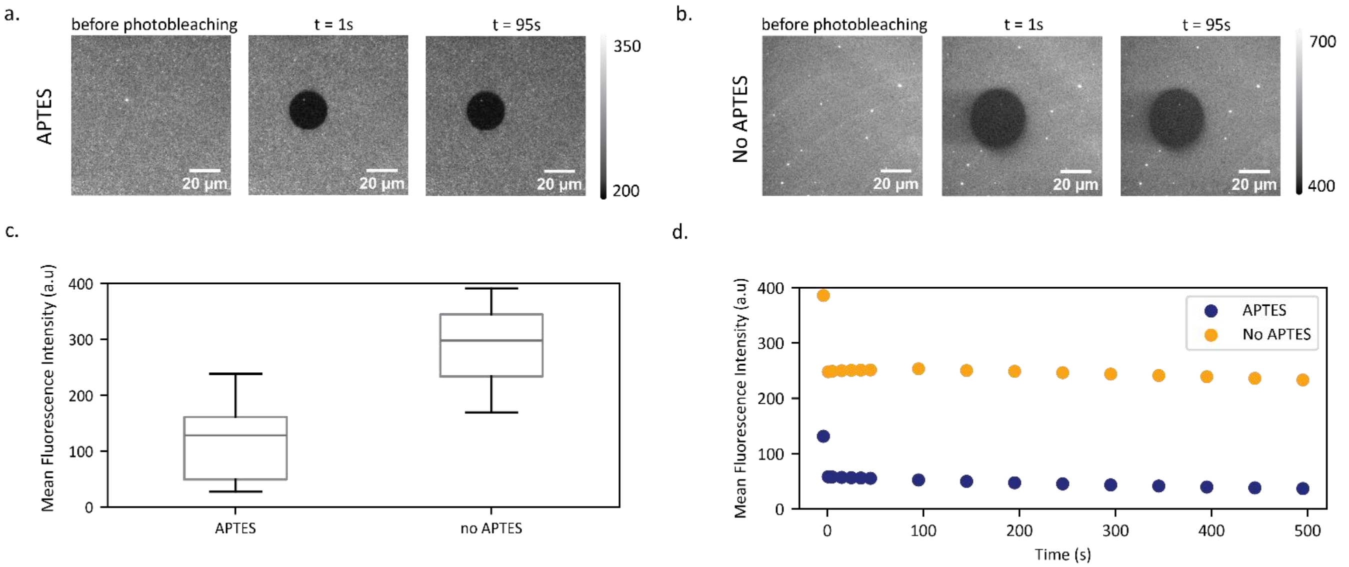

**Figure S6.** Analysis to compare the capacity APTES versus untreated glass to bind and retain NHS-PEG on the coverslip surface. a) Example frames of NHS-PEG-Cy3 on APTES surface taken before and after photobleaching (Scale bar = 20 µm). b) Same as panel (a) but PEG-Cy3 is on a surface with no APTES. c) Mean fluorescence intensity of surface with and without

APTES before photobleaching. Number of FOVs: N(APTES) = 4 and N (no APTES) = 3. d) Mean fluorescence intensity of PEG-Cy3 with and without APTES on the photobleached area of the imaged field of view (photobleaching at time = 0s). Measurements from two coverslips were averaged for each experiment.

#### S7. Single-vs. double-coating of PEG.

The protocol of Chandradoss et al.[4] describes two consecutive rounds of PEGylation. The first round contains a proportion of NHS-PEG-(*functional group*) and the second round contains only NHS-PEG with lower molecular weight to make the PEG layer denser and thereby improve the passivation.

To study the passivation quality of single-layer versus double-layer PEG coatings in the case of subsequent functionalization with DNA oligonucleotides, we prepared coverslips with single-layer PEG and double-layer PEG, both including 5 % PEG-maleimide. Control coverslips coated with PEG but without PEG-maleimide were prepared, to allow for comparison between specific and non-specific binding. In coverslip coating preparation, the first PEG layer was functionalized with maleimide, while the second layer was added for enhanced passivation of the surface.

All coverslips were then incubated with fluorescently labelled and thiol-functionalized DNA (SH-DNA-Atto633, 1.82 µM) and subsequently washed (see Methods and Materials). We imaged different coverslips for each condition in different FOVs and analyzed the fluorescence intensity.

The controls (without PEG-Mal) demonstrate the passivating effect of double PEG coating, with the non-specific binding slightly decreased compared to single-coating. This is consistent with the observations from the original protocol.

In the case of single-coated PEG-Mal surfaces, we noticed a significant increase in fluorescence intensity compared to the control confirming specifically bound DNA. In the case of double-coated PEG-Mal surface, we observed a small increase in fluorescence intensity compared to the control DNA for the PEG-Mal sample. This was unexpected, since we thought that the second PEG layer would increase passivation without interfering with functionalization. We suspect that the second round of PEGylation affects the functionality and/or the accessibility of the maleimide group, either sterically or by possibly hydrolyzing the first PEG layer during the long overnight incubation period of the second PEG layer. Further investigation would be needed.

This result led us to choose single-coated PEG surfaces in our coverslip functionalization assays.

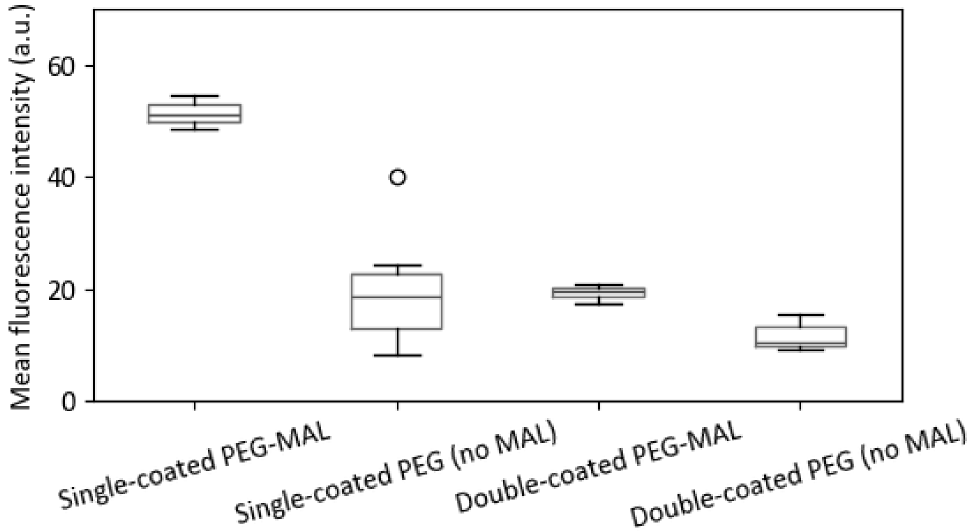

**Figure S7.** Mean fluorescence intensity of SH-DNA-Atto633 on single- vs. double-coated PEG. Single-coated PEG-Mal (5 % PEG-Mal), single-coated PEG (no Mal) (100 % PEG), double-coated PEG-Mal (5 % PEG-Mal in first round, 100 % PEG in second round), double-coated PEG(no Mal) (100% PEG in first and second round). Each surface was measured on two coverslips with a minimum of 2 FOVs per coverslip.

#### S8. Gas-phase- versus liquid-deposition of APTES.

Different protocols using vapor phase or liquid phase deposition of APTES have been reported[7]. We compared liquid- and gas-deposition methods of APTES on borosilicate glass coverslips.

We incubated fluorescent PEG (PEG-Cy3, 10%) with APTES-coated coverslips.

The sample with gas-deposited APTES shows a higher level of uniformity than the one of liquid-deposited APTES (Figure S8a). This could be due to the greater control over the humidity levels in gas-deposition, since humidity levels affect the self-polymerization of silanes[8].

To test the stability of the PEG surfaces, we conducted a photobleaching experiment (see Methods). We observed no recovery after photobleaching across an 8-minute time period (1 fps) for both APTES preparations. This demonstrates the stability of the bonds between APTES and NHS-PEG, independent of the APTES-deposition method.

Comparing the fluorescence intensity from both conditions shows liquid-deposited APTES has a higher mean value. We interpret this as a higher presence of bound PEG molecules on the liquid-deposited APTES surface coating compared to the gas-deposited counterpart. To understand the influence of the bright spots observed on liquid-deposited APTES samples, we analyzed the intensity in areas without the bright PEG spots and the same trend was observed. We hypothesize that the liquid-deposition of APTES may lead to surface defects which can bind a higher concentration of PEG.

In a control group where no APTES was used, the fluorescent PEG attached directly to the cleaned coverslip. Interestingly, in this case, an increase in fluorescence intensity was observed compared to both liquid and gas-deposited APTES coverslip surfaces and showed a spotty surface, similar to liquid-deposited APTES, but smaller spots.

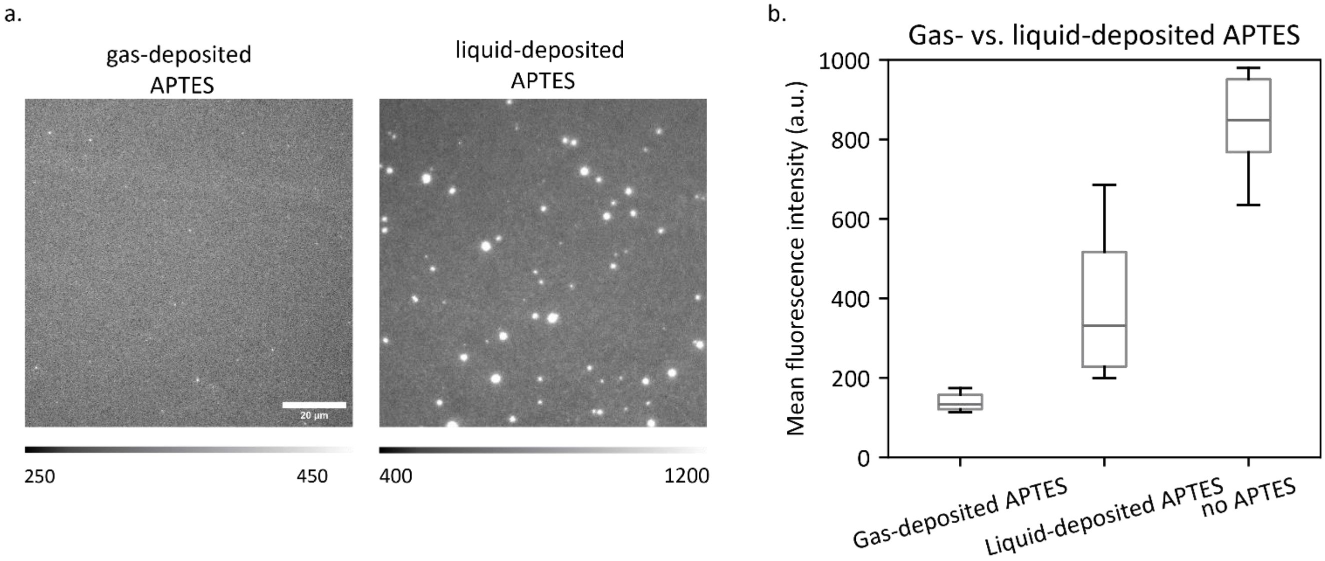

**Figure S8.** Comparison of liquid- and gas-deposition of APTES. a) Example images of gas-deposited and liquid- deposited APTES after incubation with PEG-Cy3 (Scale bar = 20 µm). b) Mean fluorescence intensity of PEG-Cy3 on gas-deposited APTES, liquid-deposited APTES and glass (no APTES). Total sample counts: each condition was tested on 2 different coverslips with each 10 field of views imaged.

### S9. Histograms of events detected on PLL-coated coverslip.

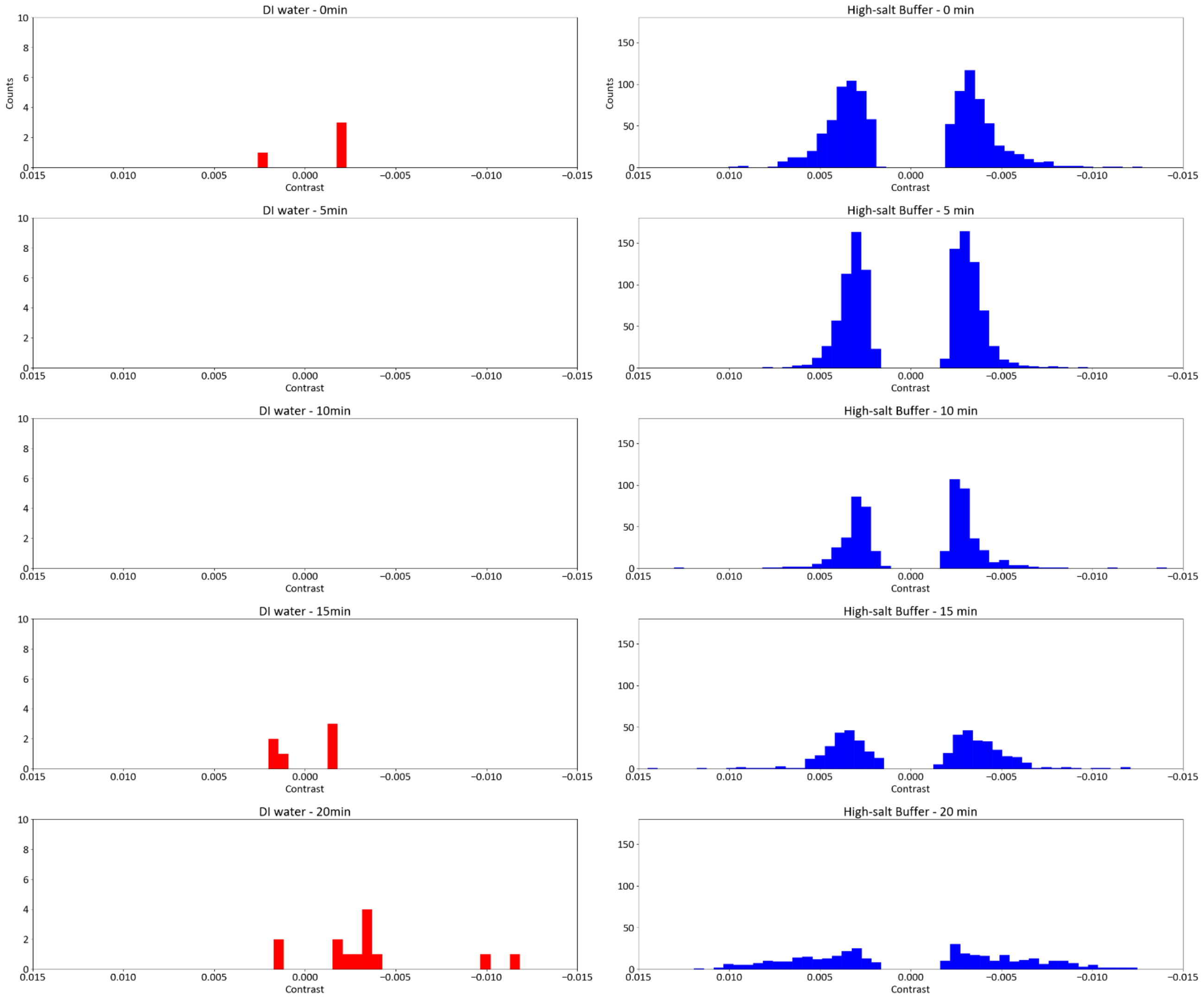

**Figure S9**. Histograms of events detected on PLL-coated coverslips with DI water and high-salt buffer measured at different times after incubation with respective buffer. This figure is related to data from Figure 3C – one example histogram for each condition is shown. Total event counts: N (DI water – 0 min) = 4, N (DI water – 5 min) = 0, N (DI water – 10 min) = 0, N (DI water – 15 min) = 6, N (DI water – 20 min) = 14, N (high-salt buffer – 0 min) = 997, N (high-salt buffer – 5 min) = 1087, N (high-salt buffer – 10 min) = 585, N (high-salt buffer – 15 min) = 479, N (high-salt buffer – 20 min) = 394.

#### S10. Photobleaching experiment for bond between PEG-Mal and Thiol-DNA.

To examine the stability of the thioether bond between the PEG-Mal and Thiol-DNA (fluorescently labelled), an area within the field of view (FOV) was photobleached and imaged over an 8-minute duration (1 fps). Notably, no recovery after photobleaching was observed in the measurements. This indicates that fluorescently labelled DNA molecules exhibited minimal mobility or were entirely immobile during the acquisition time.

This indicates that the washing steps were adequate in removing any weakly bound or excess molecules, the surface was sufficiently passivated to prevent further binding, and that the bond between PEG and DNA is strong enough under solvent incubation.

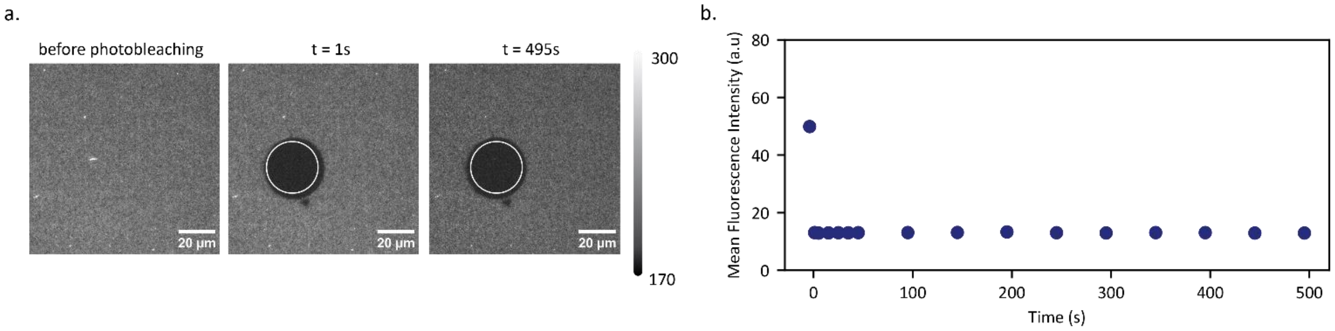

**Figure S10.** Photobleaching experiment with PEG-Mal and SH-DNA-Atto633. a) Examples of raw data of SH-DNA-Atto633 taken at different time points, before and after the photobleaching event (Scale bar= 20μm). b) Mean fluorescence intensity data of SH-DNA-Atto633 on the photobleached area of the imaged field of view (photobleaching at time= 0s). Two coverslips were imaged and show the same trend, but only one is shown.

#### S11. iSCAT measurement of fully functionalized PEG surface (APTES, SVA-PEG2K-MAL, SH-DNA)

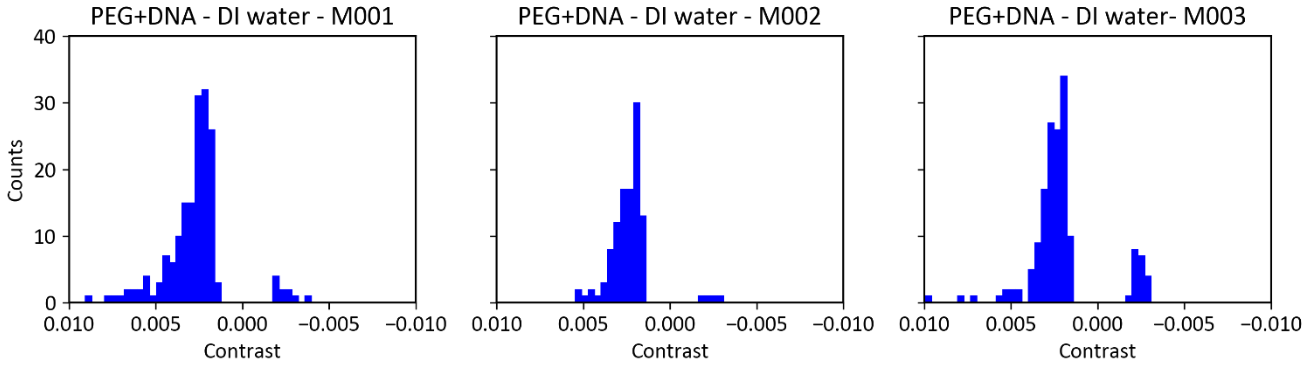

**Figure S11**. Histograms of events detected on fully functionalized PEG surfaces with DI water from 3 different coverslips. This figure is related to data from Figure 3E. Total event counts: N (M001) = 174, N (M002) = 110, N (003) = 159.

#### S12. Root-mean-square of pixel values for iSCAT data

| Dataset | RMS of individual measurements | Related figure |
| --- | --- | --- |
| PEG 30k, with DI water on iSCAT | 0.00138, 0.00115, 0.00102 | 1B,C; S1 |
| PEG 30k, with high salt buffer on iSCAT | 0.000588, 0.000600, 0.000705 | 1B,C; S1 |
| PEG 2k, with DI water on iSCAT | 0.000535, 0.000532, 0.000572 | 1E, F |
| PEG 2k, with high salt buffer on iSCAT | 0.000565, 0.000533, 0.000519 | 1E,F |
| PEG 2k + SH-DNA, with DI water on iSCAT | 0.000508, 0.000473, 0.000510 | 3E; S11 |
| PLL, with DI water on iSCAT(t=0min) | 0.000507 | 3B,C; S9 |
| PLL, with high salt buffer on iSCAT(t=0min) | 0.00119 | 3B,C; S9 |
| Uncoated glass, with DI water on iSCAT | 0.000516 | S13-2 |
| Uncoated glass, with high salt buffer on iSCAT | 0.000542 | S13-2 |
| APTES, with DI water on iSCAT | 0.000467 | S13-2 |
| APTES, with high salt buffer on iSCAT | 0.000579 | S13-2 |

**Table S12.** Root-mean-square (RMS) of pixel values for different measurements. For each calculation of the RMS 2000 frames were considered.

#### S13. Functionalization with 3-aminopropyltriethoxysilane (APTES) and Glutaraldehyde (GA) with Amine-DNA

To avoid surface functionalization based on electrostatic interactions, we investigated covalent bonding approaches involving small molecules, aiming to reducing the scattering signal of the coating and achieve better control of molecular display. Strategies exist to link molecules covalently to a surface[9], including using the linkage between amine-groups and aldehydes, e.g. in oligonucleotide- and protein-based microarrays[10,11]. Our approach includes APTES as it can bind to glass via its hydroxyl functional groups and to a wide range of molecules via its primary amine functional group[12]. We avoid a highly reactive functional group on the DNA as it could degrade rapidly when stored even for short time periods. Therefore, our strategy is to use amine-functionalized DNA (amine-DNA) and link it via glutaraldehyde (GA) to APTES (Figure S13-1).

To test the suitability of this approach for iSCAT microscopy, we measured the scattering signal of the APTES- coated coverslips with iSCAT. The overall signal and number of events detected on the functionalized coverslips were comparable to clean coverslips with DI water and high-salt buffer, respectively (Figure S13-2).

Next, we tested the ability to functionalize the APTES+GA coverslip with amine-DNA. We titrated the concentration of fluorescently-labelled amine-DNA origami and imaged the surface after washing (see SI). We noticed that the fluorescent signal appeared non-uniform (Figure S13-3), but it is unclear whether the origin is aggregated DNA origami and therefore inhomogeneous distribution of DNA, or non-uniformity of the underlying surface coating. Regarding the titration, while we observed an increase in fluorescence intensity when incubating with increasing concentration of amine-DNA, we also noticed a high signal in the control (DNA origami without the amine group). We suspect that this is due to insufficient surface coverage of the passivation layer. A portion of the surface may be positively charged from the APTES, even after the incubation with GA, allowing negatively charged DNA molecules to also bind non-specifically via electrostatic interactions.

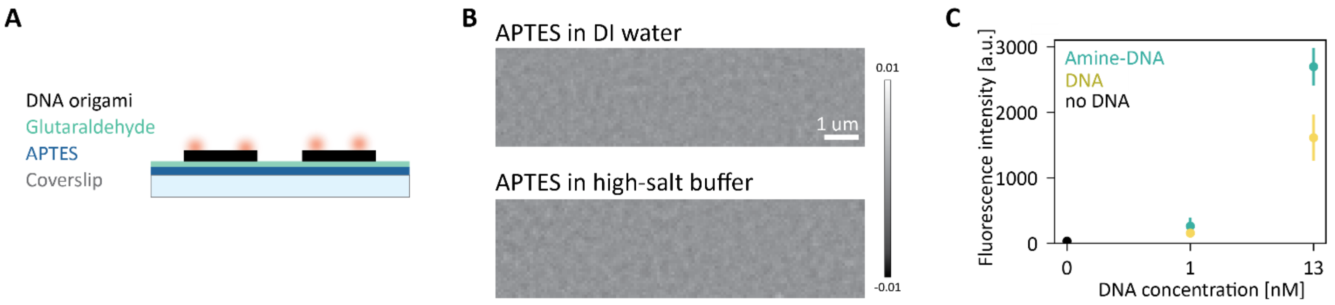

**Figure S13-1: Surface functionalization with APTES and GA.** (A) Sketch of APTES- and GA-coated coverslip with fluorescently- labelled Amine-DNA. (B) Examples of ratiometric iSCAT data of an APTES- coated coverslip incubated with different buffers. (C) Mean fluorescence intensity per FOV of APTES- and GA- coated coverslips incubated with Amine-DNA-Cy5 and DNA-Cy5. Number of measured samples: N (0 nM) = 2, N (1 nM, 13 nM) = 3.

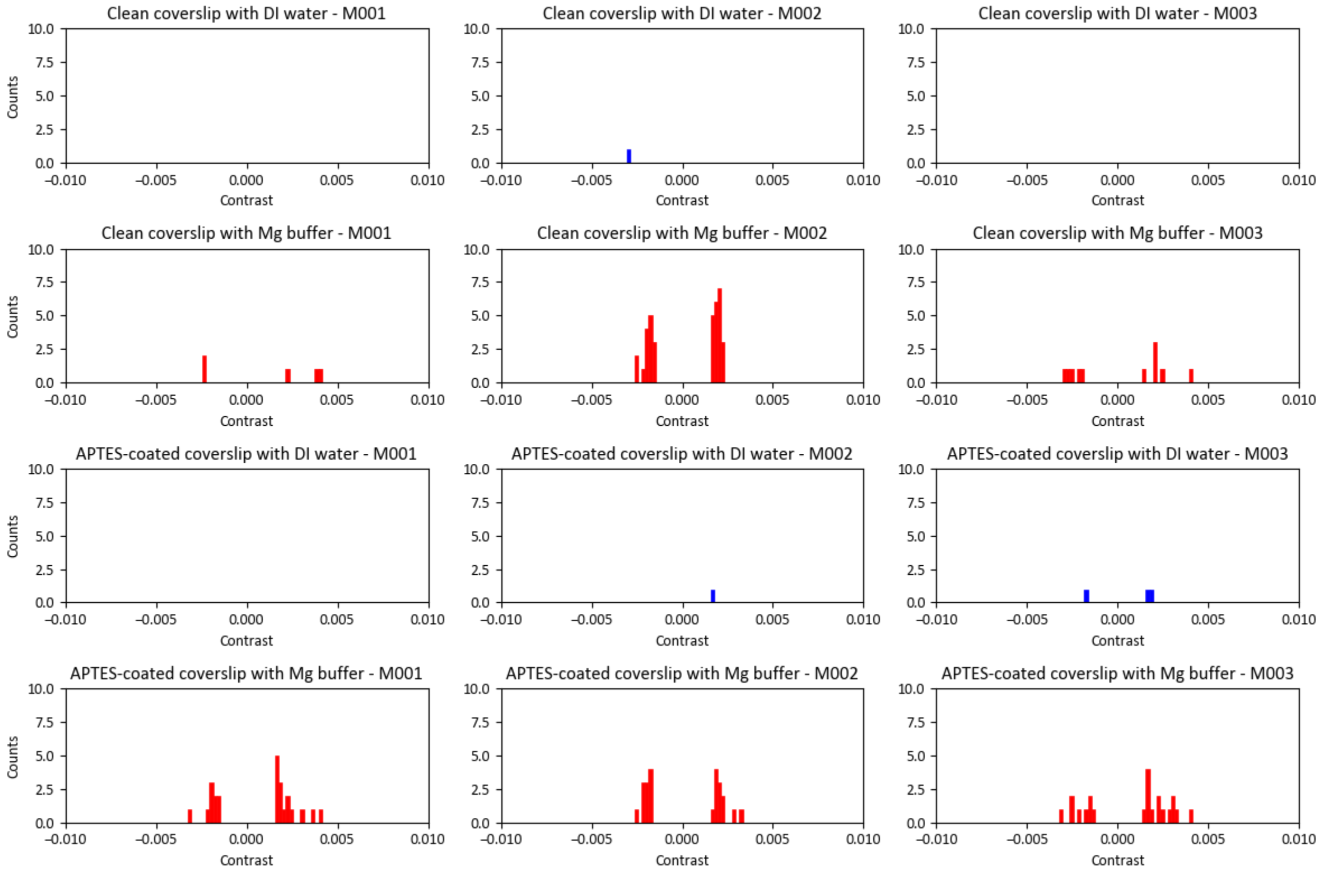

**Figure S13-2.** Histograms of events detected on clean coverslips and APTES-coated coverslips with DI water and high-salt buffer (named "Mg buffer" in the figure). Data is related to Figure S13-1B. Total event counts: N (Clean coverslip - DI water, M001, M002, M003) = 0, 2, 0; N (Clean coverslip – high-salt buffer, M001, M002, M003) = 13, 37, 12; N (APTES-coated coverslip- DI water, M001, M002, M003) = 0, 2, 4; N (APTES-coated coverslip – high-salt buffer, M001, M002, M003) = 33, 30, 99.

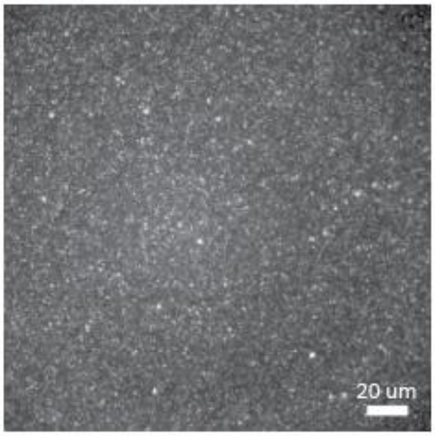

**Figure S13-3.** Cy5-labelled amine-functionalized DNA origami (13 nM) on APTES- and GA-coated surface. This figure relates to the data from Figure S13-1.
